# EZH2-driven immune evasion at disease presentation defines a targetable high-risk subset of acute leukemia exemplified by t(16;21) FUS::ERG AML

**DOI:** 10.1101/2024.05.14.594150

**Authors:** Nathaniel J Buteyn, Samantha J LaMantia, Connor G Burke, Vincent J Sartori, Eve Deering-Gardner, Zachary J DeBruine, Dahlya Kamarudin, Darrell P Chandler, Alexander C Monovich, Monika W Perez, Joanna S Yi, Rhonda E Ries, Todd A Alonzo, Russell JH Ryan, Soheil Meshinchi, Timothy J Triche

## Abstract

The past 25 years of clinical trials have produced few improvements in pediatric AML (pAML) outcomes. This is acutely evident in patients with t(16;21)(p11;q22), yielding *FUS::ERG*. Patients with *FUS::ERG*-positive AML relapse quickly and do not respond to transplantation. Major histocompatibility complex (MHC) class I & II receptors and costimulatory molecules are absent at diagnosis in *FUS::ERG-*positive AML, mirroring the phenotype and outcomes of post-transplant relapse. We show that this is driven by overexpression of *EZH2, in vitro* and in multiple clinical cohorts. While *FUS::ERG* AML is the most extreme example, this phenotype is shared by lethal *CBFA2T3::GLIS2-*driven AML, and patients with *RUNX1::RUNX1T1* have significantly worse outcomes when EZH2 overexpression co-occurs. The FDA-approved EZH2 inhibitor tazemetostat reverses this phenotype, re-establishes MHC presentation, and elicits immune effector cell-mediated elimination. EZH2 inhibitors may provide the first targeted therapeutic frontline option for AML patients with *FUS::ERG*, with the potential for broader frontline immunostimulatory benefits.

**STATEMENT OF SIGNIFICANCE:** Here we show an immune-evasive phenotype, present at diagnosis and characterized by elevated *EZH2* levels and loss of MHC class I and II, defines a high-risk subtype of acute leukemia. Treatment with the EZH2 inhibitor tazemetostat and IFN-γ reverses this phenotype and results in immune cell engagement and blast elimination.

## INTRODUCTION

The E26 transformation-specific (ETS) family of transcription factors regulate a host of cellular functions including differentiation, proliferation, and apoptosis. When aberrantly expressed, often through chromosomal translocations resulting in chimeric fusion proteins, ETS factors drive a variety of cancers^1–5^ including acute leukemias^6,7^. Translocations involving ETS factors have historically been considered high-risk in pediatric acute myeloid leukemia (pAML). In particular, the ETS family member ERG, when fused with the RNA-binding protein FUS in a t(16;21)(p11;q22) translocation, drives particularly poor prognosis in patients. Standard therapy has proven ineffective for *FUS::ERG* pAML; median time to relapse was <10 months, five-year overall survival was 33%, and 100% of patients who received stem cell transplant (SCT) died^8,9^. There is a desperate need for novel therapeutic options that target this high-risk subset.

Children’s Oncology Group (COG) pAML trials AAML0531 and AAML1031^8,9^ comprise the largest collection of pediatric AML patients with ETS fusions ever assembled. Analysis of this cohort allowed us, for the first time, to identify a pervasive immune-evasion phenotype ubiquitously present across *FUS::ERG* pAML. Through aberrant ERG activity, *FUS::ERG* pAML significantly overexpresses the histone-lysine methyltransferase EZH2 resulting in silencing of *CIITA* and downregulation of class I and II major histocompatibility complex (MHC) receptors; loss of MHC expression has been shown to strongly correlate with relapse in leukemic disease^10–13^. We demonstrate here that treatment with the FDA-approved EZH2 inhibitor tazemetostat, in combination with IFN-γ, successfully reverses the immune-evasion phenotype, decreasing blast viability in *FUS::ERG* AML cells and eliciting donor immune cell cytotoxicity. Further analysis revealed the pervasiveness of this phenotype. In the most common subtypes of pAML – those with no detectable fusion (NONE) and a t(8;21) translocation (RUNX1::RUNX1T1) – EZH2HIGH patients exhibit significantly lower MHC expression and increased rates of relapse. Immune evasion often develops in a post-relapse setting^14^, but presents here at diagnosis, providing a unique therapeutic opportunity. Targeted EZH2 inhibition from the onset of treatment may prove pivotal in restoring immunogenicity and bridging patients to SCT.

## RESULTS

### Cohort characteristics

The COG AAML0531 and AAML1031 cohorts comprise 2399 pAML patients, of which 66 patients have translocation resulting in a chimeric fusion protein containing an ETS factor. Of those 66 patients, 59 fusions contained either the ETV6 or ERG transcription factor, with *FUS::ERG* accounting for 18.2% of all ETS fusions (**Figure 1A**). The *FUS::ERG* fusions occurred in older patients (median 11.2 years) compared to *ETV6::MNX1* fusions (median 9 months). 66.7% (8/12) of *FUS::ERG* patients were female and 41.7% were Hispanic, proportions higher than what are observed across all pAML patients in AAML0531/1031 (47.9% female and 16.1% Hispanic). Of the 6 patients enrolled in AAML0531, four received gemtuzumab ozogamicin (Mylotarg) in addition to standard therapy. Five out of six AAML1031 patients were enrolled in Arm B and received bortezomib in addition to standard therapy, with one receiving standard therapy alone (Arm A)^8,9^. Taken together, 8/12 *FUS::ERG* patients achieved complete remission (CR) after the first round of chemotherapy, and 11/12 were in remission after the second round.

**Figure 1:**
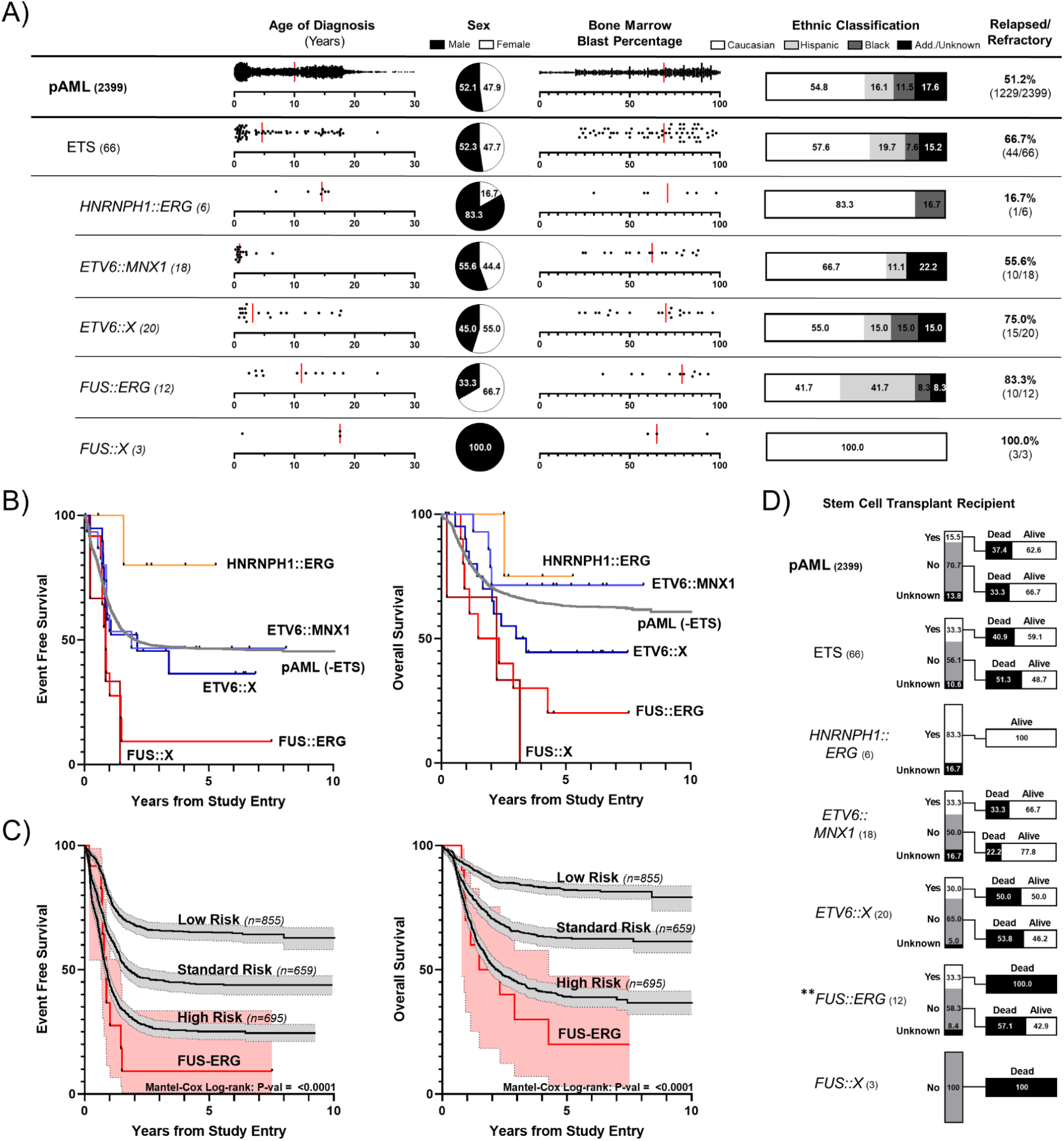
Characteristics of ETS fused pAML patients in trials AAML0531/1031. **A**) Demographic and clinical characteristics of 2399 patients enrolled in Children’s Oncology Group trials AAML0531^8^ and AAML1031^9^; major ETS fusions further expanded. **B)** Event-free and overall survival curves of ETS fused patients compared to all pAML. **C)** Survival curves for *FUS::ERG* patients compared to assigned risk categories from AAML0531/1031 (95% CI). **D)** Breakdown of frequency and success rate of hematopoietic stem cell transplant after first CR for ETS fused patients compared to all pAML. **p-val ≤0.01.

### Current therapies fail to elicit durable remission in *FUS::ERG* pAML

Overall, 51.2% of all AAML0531/1031 pAML patients had an event-free survival (EFS) event (classified as induction failure, relapse, or death), while 83% (10/12) of *FUS::ERG* patients registered EFS events (i.e., only 17% of *FUS::ERG* patients secured durable remission): 7 relapses, 2 deaths, 1 induction failure. Median EFS for *FUS::ERG* patients was 299 days (compared to 629 days for all pAML). Site of relapse/induction failure was reported for patients, excluding the two patients where primary EFS event was death, with 6/8 recorded in the bone marrow, one unknown, and one patient reporting relapse at a chloroma (myeloid sarcoma). Five-year overall survival for *FUS::ERG* patients was 33.3% (4/12), lower than the rest of pAML and other major ETS fusions, with a median time of death after diagnosis of 376 days (**Figure 1B**; **Figure 1C**). 25% of *FUS::ERG* patients (4/12) received SCT in first CR; 100% of these patients died (compared to 40.9% for ETS fusion pAML and 37.4% for all pAML) (**Figure 1D**). Given the relatively low occurrence of ETS fusion patients, it has been difficult to properly categorize the population – risk group classification from the AAML1031 protocol initially labeled 33% of *FUS::ERG* patients “low-risk”. Only in its most recent edition of “Classification of Haematolymphoid Tumors (2022)” did the World Health Organization include *FUS::ERG* as a defining genetic alteration^15^. From **Figure 1**, however, it is clear that pAML patients harboring *FUS::ERG* comprise an exceptionally high-risk population.

### EZH2-driven immune evasion

The uniquely poor prognosis for *FUS::ERG* pAML – patients often achieve a CR before rapidly relapsing and are ubiquitously nonresponsive to SCT – suggests a distinct mechanism of resistance. To evaluate this hypothesis, we analyzed RNA-sequencing data from 1,410 AAML0531/1031 patients at diagnosis, ten of whom had a *FUS::ERG* fusion. We used non-negative matrix factorization (NMF) to decompose the data into 90 factors (weighted, non-negative combinations of transcript abundances), a rank which recovered nearly all signal in the original data while simultaneously suppressing noise. The selection of features and weights that comprise each factor is data-driven, rather than pathway– or annotation-driven; each factor typically captures a discrete “aspect” of the overall data. UMAP dimension reduction on the NMF-decomposed mRNA abundance highlighted a transcriptional profile separating *FUS::ERG* patients from other pAML patients, including those harboring other ETS fusions (e.g., *ETV6::MNX1*, *ETV6::X*, and *HNRNPH1::ERG*; **Figure 2A**; two-color plot in **Supp Figure 1**). We investigated the underlying source of this separation, identifying one factor enriched for up-regulated genes, and three factors enriched for down-regulated genes (**Figure 2B**). Gene set enrichment analysis (GSEA) on the weights comprising each factor showed that some of the most down-regulated genes were immune-related, especially in NMF factor 90 (**Figure 2C**). By contrast, factor 85 (upregulated in *FUS::ERG* pAML) was driven by *EZH2, GATA2, CSF3R, IGF2BP1,* and *STAB1,* of which all save for *EZH2* have previously been reported to confer poor prognosis in pAML (**Supp Figure 2**; gene weights for all NMF factors in **Supp Table 1**). As *FUS::ERG* patients showed significant upregulation of the histone-lysine methyltransferase *EZH2* compared to all other pAML patients, including those with other ETS fusions (**Figure 2D**), we opted to investigate *EZH2*.

**Figure 2:**
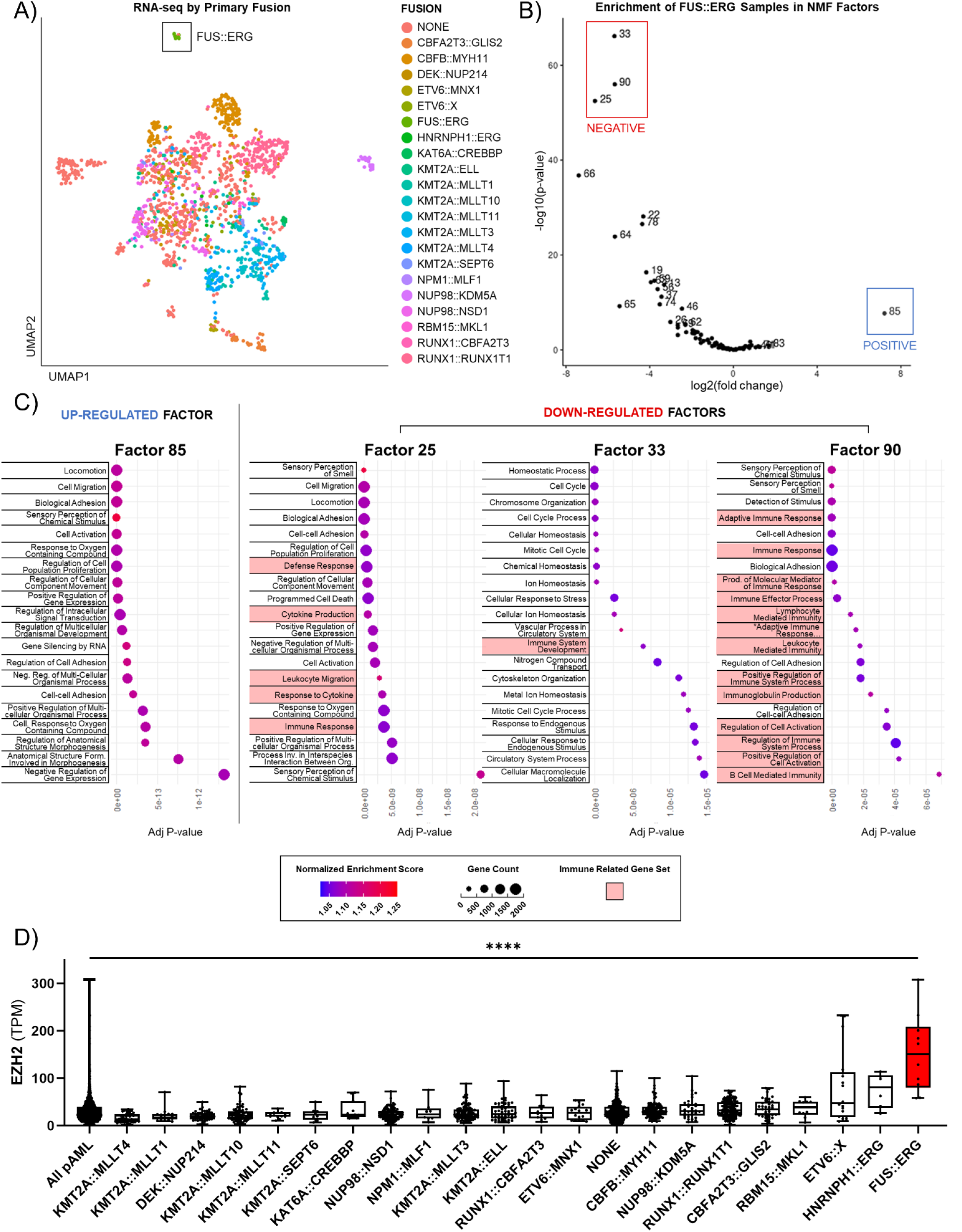
Loss of immune function defines *FUS::ERG* pAML. **A)** UMAP projection of gene expression from 1410 AAML0531/1031 patients who had RNA-sequencing at diagnosis. Sorted by major pAML fusion. **B)** Non-negative matrix factorization (90-factor) of RNA-sequencing data showing factors enriched for *FUS::ERG* signatures. **C)** Gene set enrichment analysis for Gene Ontology: Biological Processes of each enriched factor. Down-regulated factors were heavily enriched for immune related processes. **D)** Expression of *EZH2* (TPM) amongst major pAML fusions (AAML0531/1031; n = 1410). ****p-val ≤0.0001.

We then looked more closely at immune-related gene expression in *FUS::ERG* patients relative to all other pAML patients, and quantified transcript levels from the most prevalent fusion groups in AAML0531/1031 (including the four major ETS fusion subsets: *FUS::ERG*, *ETV6::MNX1*, *ETV6::X*, and *HNRNPH1::ERG*). Elevated levels of *EZH2* correlated with loss of MHC class I and II expression alongside suppression of the master regulator of MHC II, *CIITA* (**Figure 3A**). Core-binding factor AML, those patients with a t(8;21)(q22;q22) *RUNX1::RUNX1T1* or inv(16)(p13q22)/t(16;16)(p13;q22) *CBFB::MYH11* fusion, as well as those with a *RUNX1::CBFA2T3* fusion, populated the opposite side of the heatmap. These fusions displayed heightened MHC expression and immunogenic tumors characterized by low *EZH2*, high *CIITA*, and successful response to therapy in AAML0531/1031, consistent with other published cohorts^16,17^. In addition to *CIITA*, there was also significantly lower expression of *NLRC5*, a regulator of MHC I expression, in *FUS::ERG* compared to the pAML population (**Figure 3B**). Expression of *NLRC5* and *CIITA* significantly correlated with their class of MHC receptor expression (MHC I and MHC II, respectively) in the pAML patients (**Figure 3C**; n=1410).

**Figure 3:**
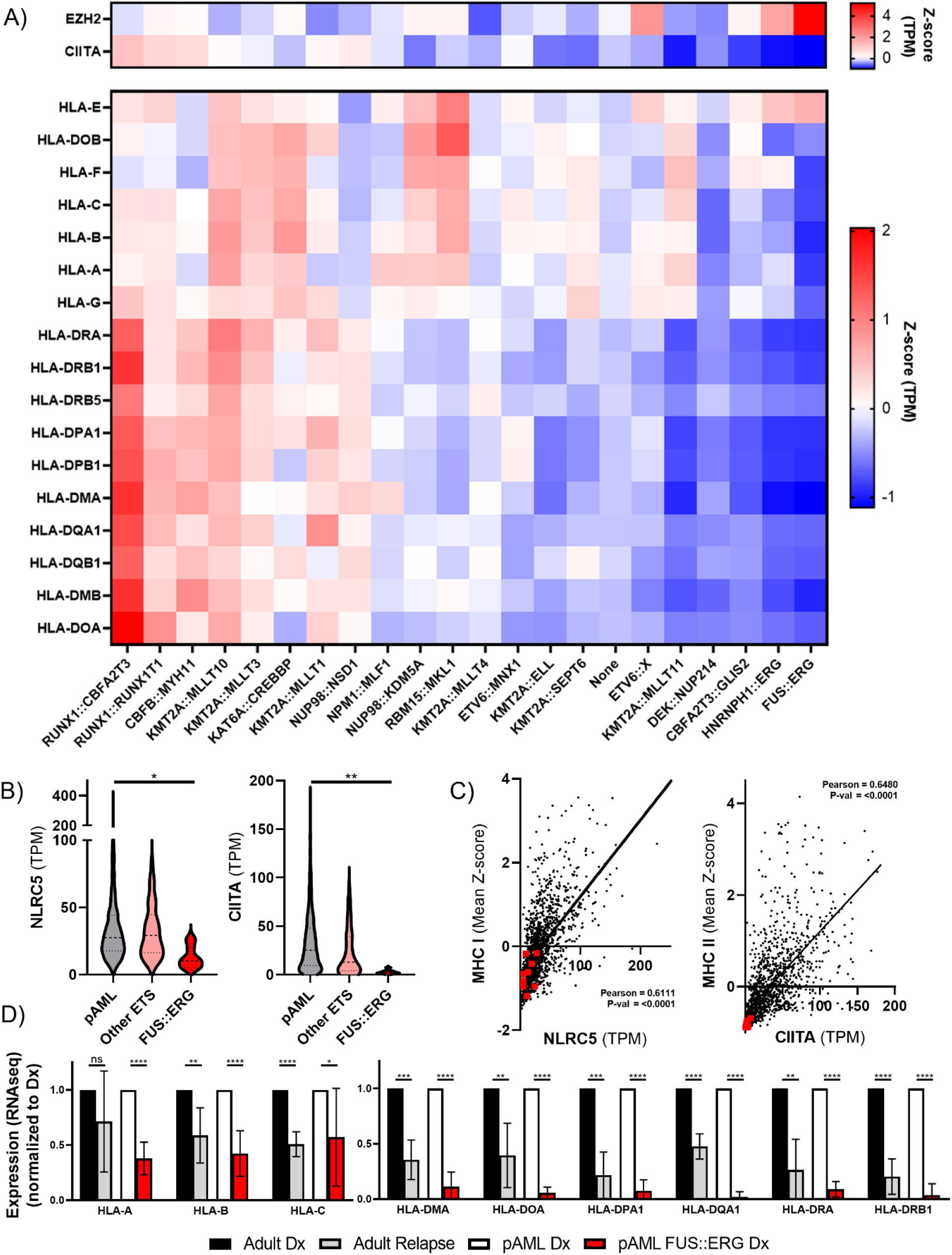
EZH2 overexpression correlates with global loss of MHC receptors. **A)** Heatmap of *EZH2*, *CIITA*, and MHC receptor expression among the major pAML fusions (TPM z-score; n = 1410). **B)** Expression of *NLRC5* and *CIITA* between *FUS::ERG* pAML, other ETS fused pAML, and the general pAML population in AAML0531/1031 (n = 1410). **C)** Scatterplot of MHC I expression (mean z-score) versus *NLRC5* in AAML0531/1031 (n = 1410; Pearson 0.8111; p-val = <0.0001) and MHC II expression (mean z-score) verses *CIITA* in AAML0531/1031 (n = 1410; Pearson = 0.6480; p-val = <0.0001); *FUS::ERG* patients highlighted in red. **D)** Expression of MHC I and II genes in adult AML at diagnosis (black) and relapse post-SCT (grey) from Christopher, *et al*.^14^ was compared to AAML0531/1031 pAML at diagnosis (white) and *FUS::ERG* pAML at diagnosis (red). ****p-val ≤0.0001; ***p-val ≤0.001; **p-val ≤0.01; *p-val ≤0.05.

Christopher, *et al.* showed that adult AML patients who relapse post-SCT demonstrate global loss of MHC receptors compared to levels at diagnosis (Adult Dx vs. Adult Relapse in **Figure 3D**)^14^. Importantly, when compared to the pAML diagnosis mean, *FUS::ERG* patients at diagnosis display the same, if not more exacerbated, MHC silenced phenotype as post-SCT relapse patients (pAML Dx vs. *FUS::ERG* Dx in **Figure 3D**). Indeed, instead of acquiring immune deficiency via the selective pressures of standard induction and SCT, *FUS::ERG* patients enter treatment overwhelmingly defined by a treatment-resistant phenotype. It is no surprise then that time-to-event is one of the shortest and response to SCT, nonexistent, compared to the rest of pAML.

AML is generally considered an immunogenic malignancy, where allogeneic SCT is effective at promoting graft-versus-leukemia (GvL) in lieu of graft-versus-host disease (GvHD)^18,19^. Effector immune cells engage their targets via MHC receptors, but also rely on a host of co-stimulatory receptors to initiate activation and facilitate clearance (**Figure 4A**). We therefore analyzed RNA-seq data for expression of ligands for natural killer (NK) cell and T cell co-stimulatory immune receptors that are typically expressed on monocytic cells. As shown in **Figure 4B**, *FUS::ERG* pAML patients showed the lowest mean expression of these ligands compared to the other major pAML fusions. Of note, the ligands for the T cell receptor CD28 (CD80, CD86) were almost completely absent in *FUS::ERG* patients. Moreover, *FUS::ERG* patients showed a significant increase in transcripts for TGF-β (*TGFB1*) and the TIM-3 ligand Galectin-9 (*LGALS9*), both inducers of immunosuppressive T regulatory (Treg) cells (**Supp Figure 3**)^20–22^. The resistance of *FUS::ERG* patients to standard therapy might therefore be explained by both the existence of an immune-desensitized leukemic blast population and a stunted immune effector cell population.

**Figure 4:**
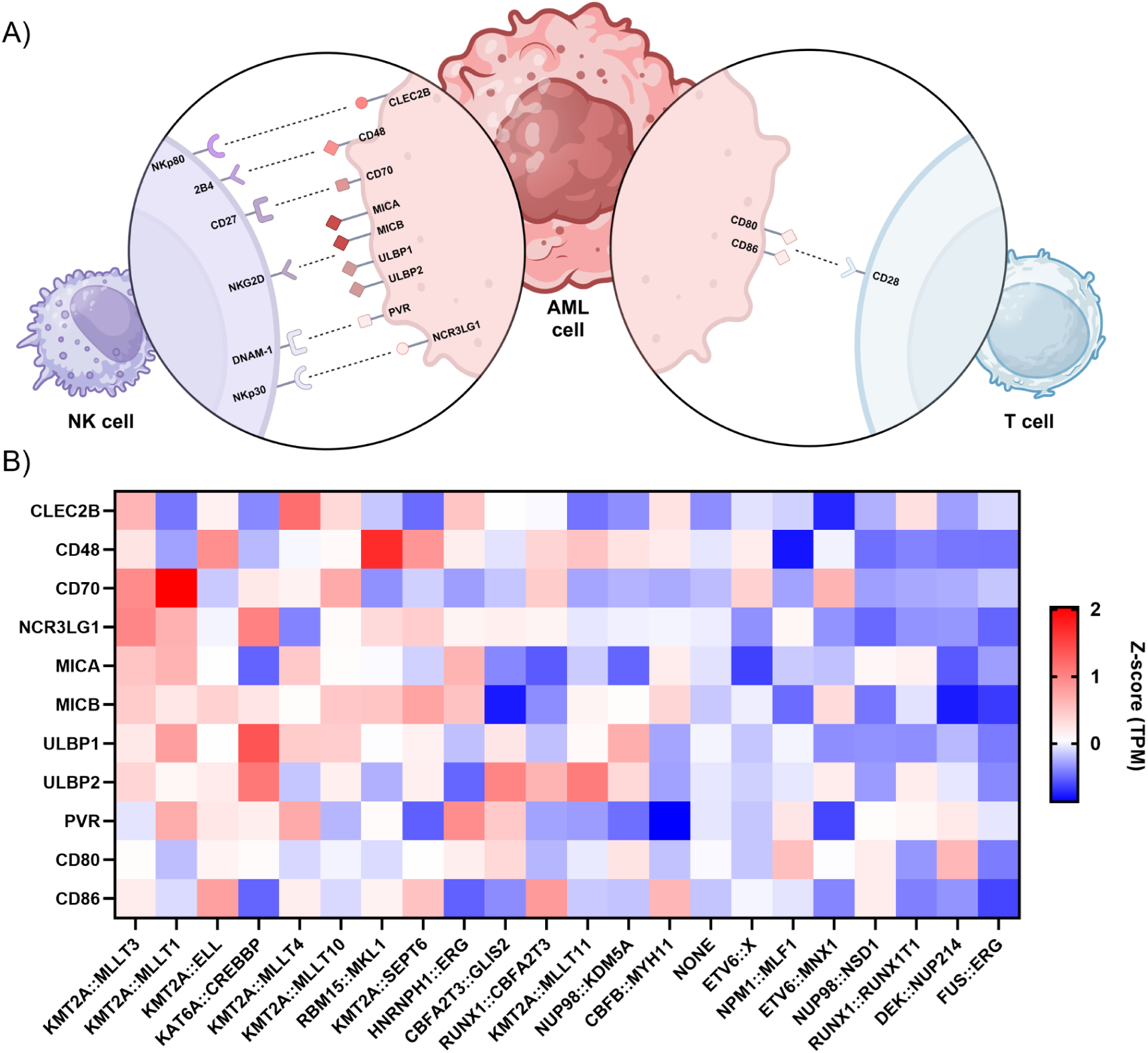
Immune co-stimulatory receptors are downregulated in *FUS::ERG*. **A)** Natural killer cell and T cell immune receptor ligands found on the surface of pAML blasts. Graphic created with biorender.com. **B)** Expression of ligands in the major pAML fusions (TPM z-score; n = 1410).

### FUS::ERG drives EZH2 over-expression

All ETS transcription factors contain a winged helix-turn-helix DNA binding domain that interacts with the GGA(A/T) motif known as an ETS-binding sites (EBS). Numerous variants of the *FUS::ERG* fusion have been reported in pediatric leukemias, yet all retain their DNA binding domain^23^. Additionally, repeats of the GGAA sequence have been shown to recruit ETS proteins and facilitate initiation of transcription^24^. Chromatin immunoprecipitation sequencing (ChIP-seq) for ERG in the patient-derived *FUS::ERG* cell line TSU-1621MT shows binding of ERG to one such GGAA repeat site in the enhancer region of *EZH2* (**Figure 5A**; data from Sotoca, *et al.*^25^). In primary pAML patient H3K27ac ChIP-seq data provided from Texas Children’s Hospital (data from Perez, *et al.*^26^) the GGAA repeat site in EZH2 was heavily acetylated, indicative of higher transcriptional activation, in only the *FUS::ERG* patient. Other patients, including those with MLL fusions, core-binding factor AML, and even another ETS fusion – *ETV6::MNX1* – showed no H3K27ac at the locus (**Figure 5B**). In fact, the only cell line that matched the H3K27ac profile of *FUS::ERG* was a Ewing’s Sarcoma line containing a *EWSR1::ERG* fusion (CHLA25; **Supp Figure 4**). Further analysis of ERG-driven cell lines demonstrated that ERG in *FUS::ERG* pAML uniquely bound both the promoter and enhancer of EZH2, suggesting the chimeric fusion may possess unique neomorphic binding affinities – even when compared to other ERG fusions such as *EWSR1::ERG* Ewing’s Sarcoma (**Supp Figure 4**). Predictably, and matching the RNA-seq data from AAML0531/1031, the *FUS::ERG* patient additionally showed a lack of accessible chromatin marks at HLA loci – including HLA-DRA shown in **Figure 5C**. Interestingly, further analysis of H3K27ac at EBS GGAA repeat enhancers across pediatric malignancies revealed that *FUS::ERG* expressed a global enrichment of acetylation at 6x, but not 3x, sites, unique amongst the other pediatric leukemias and similar only to the ETS-driven Ewing’s Sarcoma (**Figure 5D; Supp Figure 5**).

**Figure 5:**
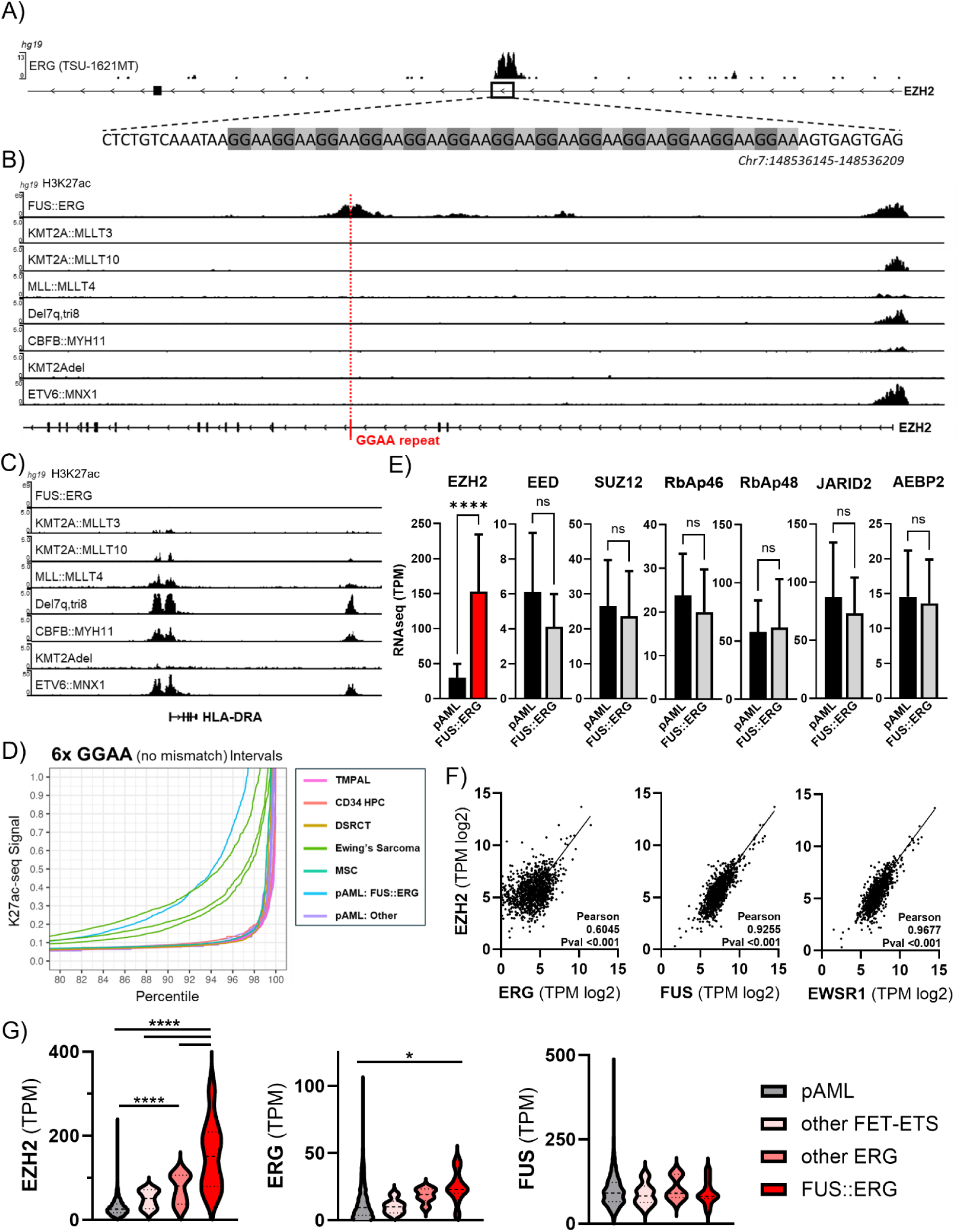
Aberrant FUS::ERG activity correlates with overexpression of *EZH2*. **A)** ChIP-seq for ERG in TSU-1621MT *FUS::ERG* cells at the GGAA repeat sequence in the enhancer region of *EZH2*^25^. **B,C)** H3K27ac ChIP-seq in primary pAML patients^26^ at the EZH2 (B) and HLA-DRA loci (C). **D)** H3K27 acetylation analysis at 6x GGAA motifs conducted on primary pAML^26^, T-lymphoid/myeloid mixed-phenotype acute leukemia (TMPAL)^61^, Desmoplastic small round cell tumors (DSRCT)^62^, and Ewing’s Sarcoma^63^ patients with CD34 hematopoietic progenitor cells (HPC) and mesenchymal stem cell (MSC) controls^64^; pAML *FUS::ERG*^26^ cases highlighted in blue. Controls shown in Supplemental figure 4. **E)** Expression of PRC2 members in AAML0531/1031 pAML compared to *FUS::ERG* (n = 1410). **F)** Scatter plots of expression of *EZH2* versus *ERG* (Pearson = 0.6045; p-val = <0.001), *FUS* (Pearson = 0.9255; p-val = <0.001), and *EWSR1* (Pearson = 0.9677; p-val = <0.001) (n = 1410). **G)** Expression of *EZH2*, *ERG*, and *FUS* among AAML0531/1031 pAML, other FET-ETS, other *ERG*, and *FUS::ERG* (n = 1410). ****p-val ≤0.0001; *p-val ≤0.05.

EZH2 itself operates canonically as the catalytic subunit of the polycomb repressive complex 2 (PRC2) to carry out histone methyl transferase activity^27,28^. Expression of other members of the complex, both core (EED, SUZ12) and auxiliary subunits (RbAp46, RbAp48, JARID2, AEBP2), are not significantly altered in *FUS::ERG* compared to the rest of pAML (**Figure 5E**). ERG has been previously shown to regulate *EZH2* in prostate cancer, where up to fifty percent of patients harbor the *TMPRSS2::ERG* fusion, localizing to the *EZH2* promoter and inducing transcription^29^. Analysis of RNA-seq from AAML0531/1031 supported these findings in pAML – a correlation is seen between expression of *ERG* and *EZH2* (Pearson = 0.6045)(**Figure 5F**). Interestingly, there was a strong correlation observed between the expression of *FUS* and *EZH2* (Pearson = 0.9255). *FUS*, or Fused in Sarcoma, is part of the FET family of RNA-binding proteins; other members include *TAF15* and *EWSR1*. Like *FUS*, a strong correlation was observed between transcript levels of *EWSR1* and *EZH2* (Pearson = 0.9677); *TAF15* did not correlate (Pearson = –0.2095) (**Figure 5F**). As both wild-type *ERG* and *FUS* correlate with *EZH2* expression, it is unsurprising that their fusion, possibly contributing aberrant activity of both members, significantly upregulates *EZH2* expression above the general pAML population as well as above patients with a non-ERG FET-ETS fusion (e.g. *FUS::FEV*, *FUS::FLI1*, *EWSR1::ELF5*, *EWSR1::FEV*) or another –ERG fusion (e.g. *HNRNPH1::ERG*)(**Figure 5G**). Furthermore, only *FUS::ERG* showed elevated wild-type ERG expression, potentially through direct transcriptional control (an EBS exists within the *ERG* gene). Wild-type *FUS* was not differentially expressed between all pAML, other FET-ETS, other –ERG fusions, and *FUS::ERG* (**Figure 5G**). The lack of heightened *EZH2* observed in other FET-ETS fusions when compared to *FUS::ERG* suggests that aberrant FUS activity from the N-terminus chimeric protein fragment may not be sufficient to over-induce *EZH2*. The detailed mechanism behind this distinct ability of FUS::ERG pAML to robustly upregulate EZH2 – and the role that each protein in the chimeric fusion may play – warrants further investigation, but lies outside the scope of this current study.

### Tazemetostat restores MHC expression

To test if inhibition of EZH2 could reverse the immune-evasion phenotype, we treated patient-derived *FUS::ERG* cell lines (TSU-1621MT and YNH-1) with the FDA-approved EZH2 inhibitor tazemetostat. The reversal of suppressive epigenetic H3K27me3 marks by tazemetostat allows access to promoter-IV (P-IV) *CIITA,* which can in turn be driven transcriptionally through IFN-γ; it has been long established that IFN-γ potently induces *CIITA* via P-IV and results in expression of MHC II molecules^30,31^. As predicted, combined tazemetostat and IFN-γ treatment robustly synergized to significantly upregulate MHC II receptor surface expression after 7 days of treatment *in vitro* compared to untreated and single treated cells (TSU-1621MT in **Figure 6A,B**; YNH-1 in **Figure 6G**). MHC I molecules were also upregulated significantly – CIITA has shown to moderately control this class as well^32,33^. Additionally, tazemetostat significantly reduced cell viability (TSU-1621MT in **Figure 6C**; YNH-1 in **Figure 6H**). Dose response experiments with tazemetostat suggested no further upregulation of MHC II occurred *in vitro* beyond 10 µM (**Figure 6D**) while significant cell death began at 1.0 µM (**Figure 6E**). Because the cell lines used were derived from adult *FUS::ERG* patients, we confirmed that the immune-evasive phenotype was present in the cells. Flow cytometry data comparing untreated cells to unstained controls (representative histogram in **Figure 6F**) shows near global loss of MHC II on the surface of the cells. Additionally, a non-ETS control AML cell line (OCI-AML3) showed no response to the combination treatment: tazemetostat + IFN-γ failed to increase expression of MHC I and any increase in MHC II expression was due to IFN-γ alone (**Figure 6I**). To assess whether the effects we observed were due to direct inhibition of EZH2, we analyzed global H3K27me3 levels after tazemetostat treatment; complete loss was observed in the Tazemetostat and Tazemetostat + IFN-γ treatments (**Figure 6J**; representative blot shown). Next, we directly degraded EZH2 via the Proteolysis Targeting Chimera (PROTAC) MS177^34^. Together with IFN-γ, treatment with MS177 was able to significantly upregulate MHC II receptor expression on the cell surface (**Figure 6K**). To assess the functional effects of MHC re-expression on the *FUS::ERG* cells, we co-cultured treated cells with peripheral blood mononuclear cells (PBMCs) isolated from healthy donors. In the presence of PBMCs, significant reduction in live cells, indicative of immune cell elimination, was observed only in the Tazemetostat + IFN-γ treated cells (**Figure 6L**).

**Figure 6:**
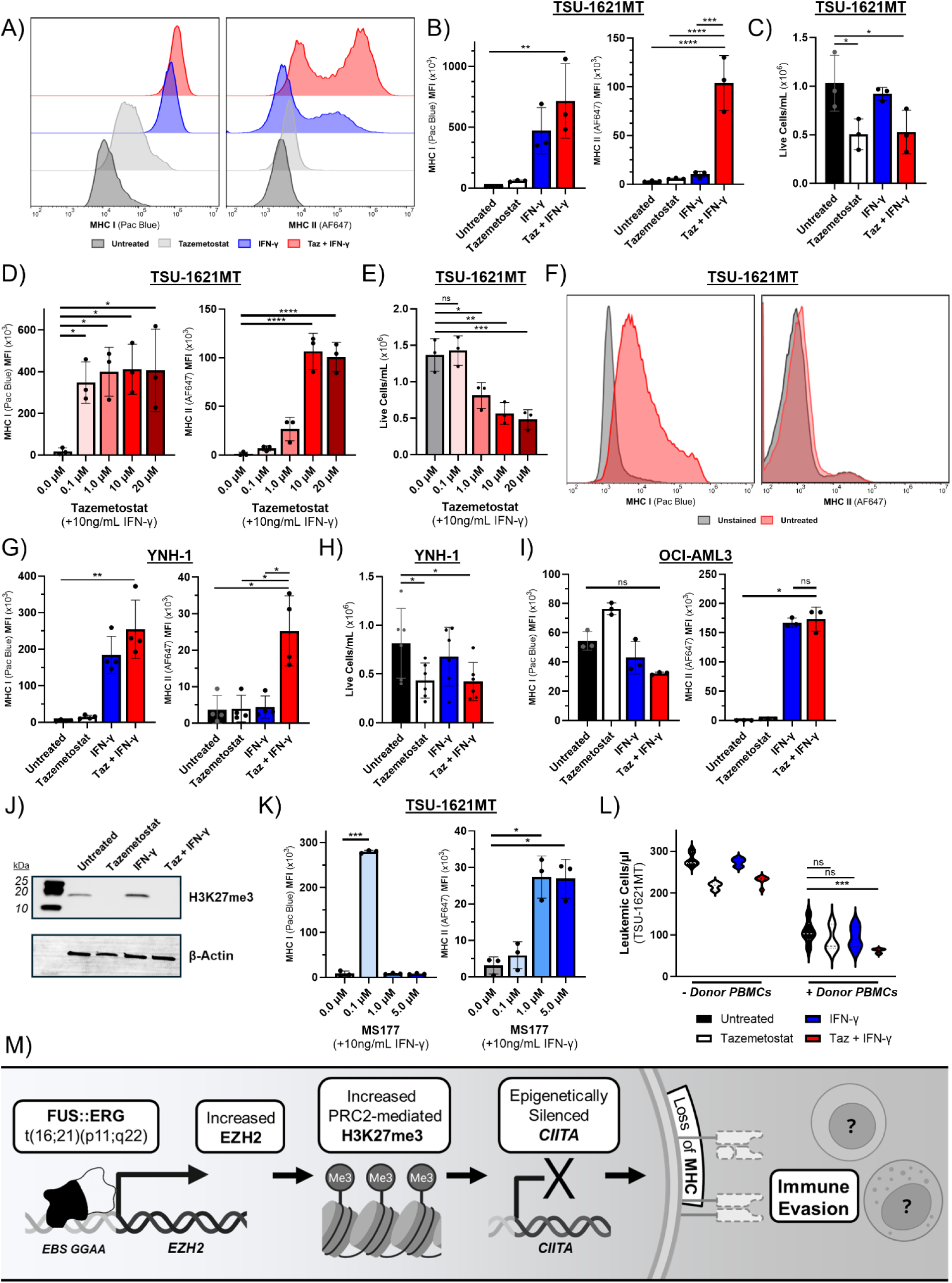
Tazemetostat restores MHC receptor expression in *FUS::ERG* cells. **A,B)** TSU-1621MT cells were treated with 10 μM tazemetostat and/or 10 ng/ml IFN-γ on days 0, 2, 4, and 6. Cells were collected on day 7 and surface expression of MHC I and II receptors was assessed via flow cytometry. Representative histogram shown in (A) and mean florescent intensity (MFI) graphed in (B) (n = 3). **C)** Cell viability in TSU-1621MT cells measured by trypan blue exclusion assay on day 6 of treatment (schedule identical to that in A for all following experiments; n = 3). **D,E)** Dose response with tazemetostat + 10 ng/ml IFN-γ. MHC flow cytometry shown in (D)(n = 3) and trypan exclusion cell viability shown in (E)(n = 3). **F)** MHC I and II flow cytometry comparing untreated and unstained TSU-1621MT cells; representative histogram shown. **G,H)** MHC I and II flow cytometry (G)(n = 4) and trypan blue exclusion cell viability assay (H)(n = 6) in YNH-1 cells. **I)** MHC I and II flow cytometry in OCI-AML3 (n = 3). **J)** Western blot for H3K27me3 in TSU-1621MT cells; β-Actin as a loading control (n = 3; representative blot shown). **K)** MHC I and II flowcytometry for TSU-1621MT treated with the PROTAC MS177 + 10 ng/ml IFN-γ; schedule identical to that in A (n = 3). **L)** TSU-1621MT cells were treated for 7 days according to the schedule described in A after which they were cocultured with PBMCs isolated from leukocyte reduction cone from healthy donors (n = 3). 20:1 ratio of PBMCs to TSU-1621MT was used and cells were cultured for 24 hours in 20 ng/mL IL-2 before cell counts were analyzed via flow cytometry. **M)** Proposed mechanism for immune suppression in *FUS::ERG* pAML. ****p-val ≤0.0001; ***p-val ≤0.001; **p-val ≤0.01; *p-val ≤0.05.

Given our experimental and clinical data, we propose that aberrant activity from the chimeric fusion protein *FUS::ERG* directly induces *EZH2* transcription. Elevated EZH2 carries out its canonical deposition of H3K27me3 which leads to silencing of *CIITA* and suppression of immune receptor expression. Unchecked by host immunity, or donor immunity in the scenario of SCT, leukemic blasts proliferate unfettered, leading to rapid relapse, SCT failure, and one of the poorest survival rates in childhood leukemia (**Figure 6M**). The robust reversal of this immune-evasion phenotype in patient-derived cells by tazemetostat suggests it may be a novel therapeutic option for *FUS::ERG* pAML patients.

### EZH2-driven immune evasion exists beyond *FUS::ERG*

To test if this phenotype existed beyond *FUS::ERG* patients, we examined the top expressors of EZH2 in the AAML0531/1031 cohort (EZH2HIGH defined here as RNAseq TPMs >1 SD above the mean) (**Figure 7A**). Mirroring what we observed in the *FUS::ERG* cases, EZH2HIGH patients showed significant down-regulation of MHC II receptors (**Figure 7B**). As expected, this group contained our *FUS::ERG* fusions but also a snapshot of the rest of the pAML cohort: other high-risk fusions like *CBFA2T3::GLIS2*, patients that had no detectable fusion (NONE), as well as the low-risk core-binding factor fusions *RUNX1::RUNX1T1* and *CBFB::MYH11*. The *FUS::ERG* and *RUNX1::RUNX1T1* fusions appeared more frequently in the EZH2HIGH group compared to AAML0531/1031 as a whole. 41.2% of patients in the EZH2HIGH classification possessed a fusion that appeared at a frequency of <5% within the group (**Figure 7C**). Interestingly, when we compared event-free survival times between EZH2HIGH and the rest of patients within the top represented fusions, we saw a significant decrease in time to adverse event for EZH2HIGH patients with no fusion (NONE) as well as a *RUNX1::RUNX1T1* fusion (**Figure 7D**). While p(16;21) *FUS::ERG* pAML consistently presents as the most severe example of this phenotype, it appears that high EZH2, irrespective of driving fusion, allows for an immune-evasive blast population that prevents sustained remission.

**Figure 7:**
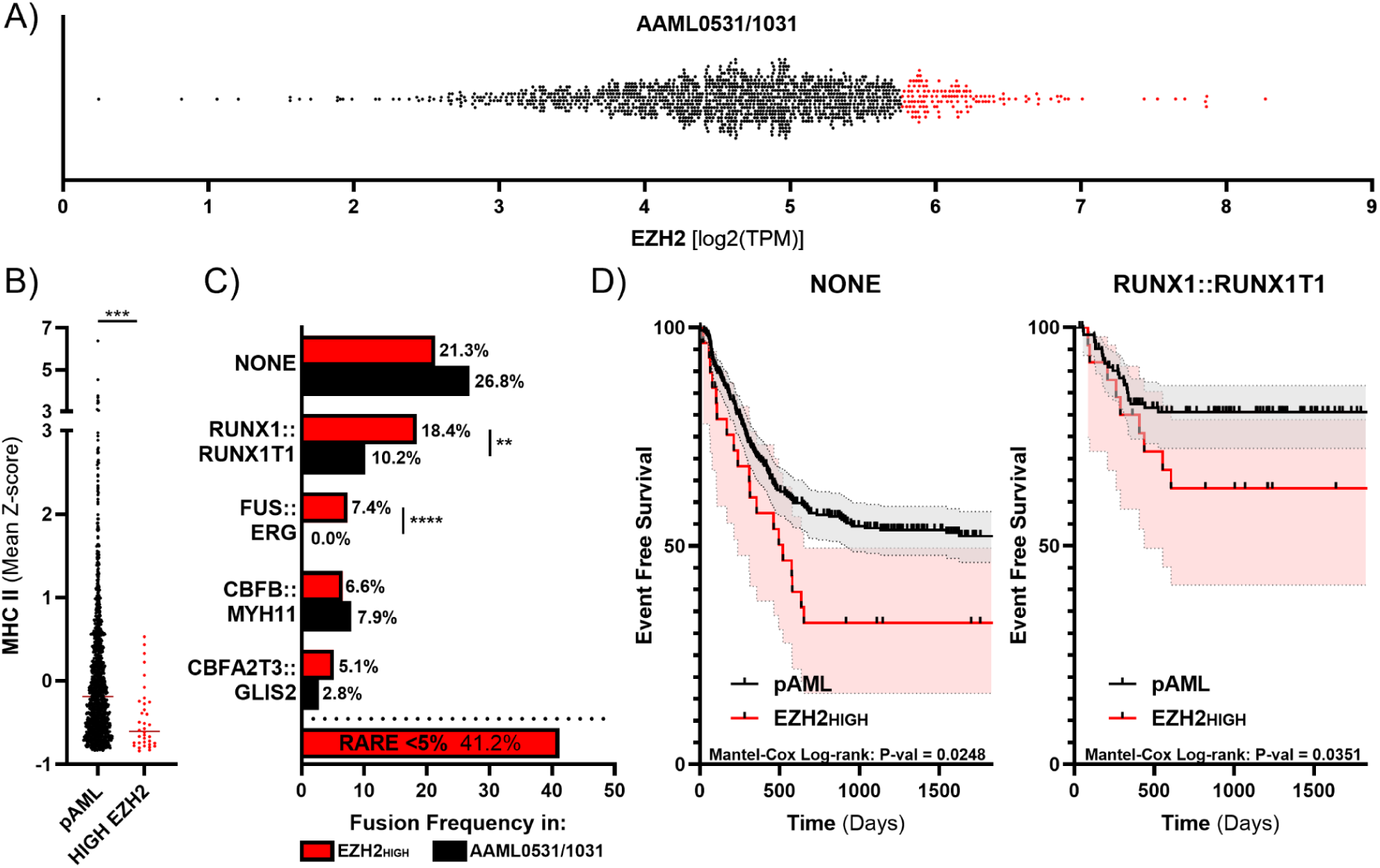
EZH2HIGH pAML exhibits similar phenotype to *FUS::ERG* patients. **A)** Patients with elevated expression of EZH2 (RNAseq TPM) within AAML0531/1031 in red and defined by TPM > 1SD above the mean (n = 136). **B)** MHC II expression mean Z-score between EZH2HIGH (n = 136) and the rest of pAML (n = 1274). **C)** Frequency of fusion type in EZH2HIGH compared to AAML0531/1031. **D)** Event-free survival curves between EZH2HIGH and the rest of pAML patients with NONE or a *RUNX1::RUNX1T1* fusion. ****p-val ≤0.0001; ***p-val ≤0.001; **p-val ≤0.01.

## DISCUSSION

The breadth of AAML0531/1031 allowed for analysis of a cohort of ETS patients the size of which was not previously available. Here, we show that *FUS::ERG* fusions drive EZH2 overexpression, a novel immune-evasion phenotype, and adverse outcomes in pAML patients. Our data and conceptual model (**Figure 6M**) are consistent with known roles for EZH2 in silencing *CIITA*, an essential component of the enhanceosome that positively regulates MHC II molecules, via P-IV chromatin H3K27me3^35,36^. The absence of MHC receptors on leukemic blasts (**Figure 3**) and the resulting loss of immunogenicity is explicitly tied to relapse in AML^10–14,37^. Monocytic leukemic blasts also express activating and inhibitory ligands on their cell surface, which interact with effector immune cells receptors to mediate immunosurveillance and target elimination. For example, NKG2D ligands increase susceptibility to allorecognition and NK cell elimination^38^; activity of NKp30 with its ligand NCR3LG1 (B7-H6) is correlated with improved overall survival in AML^39,40^; patients with high expression of DNAM-1 and its ligands have significantly longer remission and overall survival^41^; and expression of CD80 and CD86 is associated with prolonged remission^42^. AML patients with an activating NK ligand phenotype also see a significant improvement in 2-year overall survival (59.6% vs 24.4%)^43^. In *FUS::ERG* patients, these ligands were down-regulated while inducers of Treg were upregulated (**Figure 4; Supp Figure 3**). Thus, it appears *FUS::ERG* patients enter treatment at diagnosis with an immune-desensitized blast population that is not susceptible to standard induction or SCT, resulting in an overall prognosis that is one of the worst in pediatric leukemia (**Figure 1**).

Induction and intensification chemotherapies release tumor neoantigens which, when loaded on MHC receptors, elicit target elimination via effector immune cells^44^; renewal of this anti-leukemic mechanism could prove significantly beneficial in pediatric *FUS::ERG* patients. Immediate intervention at time of diagnosis could successfully bridge patients to transplant where further maintenance could elicit the desired GvL. Moreover, treatment with tazemetostat may be particularly effective in conjunction with allo-SCT to elicit potent GvL effects. There is a window immediately following SCT where endogenous inflammatory cytokines such as IFN-γ are elevated^45,46^. Treating these patients with tazemetostat should make *CIITA* transcriptionally accessible to the inevitable interferon cascade, up-regulating MHC receptors and enabling donor lymphocytes to engage with any residual leukemic blasts.

Modeling ETS fusions, and *FUS::ERG* in particular, brings certain challenges, both due to the relatively limited patient population as well as the inherent biology that defines the fusions. The locations of ETS-binding GGAA tandem repeat sites has been shown to be poorly conserved between humans and rodents^47^, and prevents accurate modeling of the ETS-driven mechanisms of leukemogenesis explored here (**Figure 5A**). Moreover, giving the vital role immunity plays in *FUS::ERG* biology, xenograft modeling in immune-incompetent mice is also ineffective. For this study we utilized patient-derived cell lines that matched the MHC II absent phenotype observed in pAML *FUS::ERG* patients (**Figure 6F**). Given the ubiquitousness of the phenotype amongst patients – all AAML0531/1031 *FUS::ERG* patients expressed EZH2 at least 1 standard deviation above the pAML population mean; mean standard score was 6.15 (**Figure 2D**) – and the robust reversal of the observed phenotype after tazemetostat treatment (**Figure 6**), consideration of EZH2 inhibition for this high-risk subset is warranted.

Given the role EZH2 plays in driving oncogenesis in a number of malignancies^48,49^, there has been considerable effort focused on developing EZH2 small molecule inhibitors. Tazemetostat (Tazverik) is an orally available, S-adenosyl methionine (SAM) competitive inhibitor of EZH2 that has been approved by the FDA for adults (16+) with metastatic or locally advanced epithelioid sarcoma and adults with relapsed or refractory follicular lymphoma^50^. Tazemetostat was reported to be well tolerated with a favorable safety profile^51,52^. It is also being used to suppress oncogenic transformation and sensitize Ewing’s sarcoma patients to CAR T cell therapy^48,53^, for treatment of pediatric rhabdoid tumors harboring SMARCB1/SMARCA4 or EZH2 alterations^54^, and is under consideration for prostate cancer patients with a TMPRSS2::ERG fusion and aberrant EZH2 activity^55^. Here we demonstrate that Tazemetostat may have even broader applicability. While *FUS::ERG* pAML is the most potent example of this immune evasion phenotype, pAML patients with high EZH2 – regardless of fusion type, risk-classification, or any other patient characteristics – have significantly suppressed immune signaling receptor expression and relapse at rates quicker than their EZH2NORMAL counterparts (**Figure 7**).

In summary, there is currently no effective treatment for *FUS::ERG* pAML. Here we have identified, for the first time, a distinct immune evasion phenotype present at diagnosis and readily targetable by the EZH2 inhibitor tazemetostat. Moreover, this phenotype appears beyond ETS fusions – even in common and/or favorable-risk pAML subtypes (NONE, *RUNX1::RUNX1T1*), EZH2HIGH patients have significantly worse outcomes. Reinvigoration of the patient’s immune response has the potential to eliminate residual disease after standard induction and elicit potent graft-versus-leukemic effects upon the introduction of donor immune effector cells (via SCT or donor lymphocyte infusions). Successful translation of these results may hold the key to improving clinical outcomes in EZH2HIGH acute leukemia.

## DATA AVAILABILITY

The data that support the findings of this study are available via dbGaP (RRID:SCR_002709) upon authorized access request, with associated metadata organized and available from the Genomic Data Commons portal (RRID:SCR_014514). The accessions include TARGET: Acute Myeloid Leukemia (AML), phs000465; Gabriella Miller Kids First! Pediatric Research Program in Germline and Somatic Variants in Myeloid Malignancies in Children, phs002187; TARGET: Acute Lymphoblastic Leukemia (ALL), phs000463, phs000464 & phs000218; TCGA LAML, phs000178; and BEAT-AML (Functional Genomic Landscape of Acute Myeloid Leukemia), phs001657.

## ACKNOWLEDGEMENTS

Research was supported by Van Andel Institute’s T32 Cancer Epigenetics Training Program (5T32CA251066-03), the Children’s Oncology Group, the National Cancer Institute (NCTN Operations Center Grant U10CA180886, NCTN Statistics & Data Center Grant U10CA180899, and R03CA290259 to TJT), the National Institute for Allergy and Infectious Diseases (R01AI171984 to TJT), the Michelle Lunn Hope Foundation, St. Baldrick’s Foundation, and by COG clinical trial participants and their families.

Disclaimer: The content is solely the responsibility of the authors and does not necessarily represent the official views of the National Institutes of Health.

## METHODS

### AML Cohort

Patient data from the COG trials AAML0531^8^ (NCT01407757) and AAML1031^9^ (NCT01371981) were used for this study. AAML0531 inclusion criteria extended to all AML patients up to 29 years of age. AAML1031 inclusion criteria extended to newly diagnosed or untreated primary AML patients up to 29 years of age, with some additional restrictions on blast count and pathognomonic cytogenetic findings. Exclusion criteria for AAML1031 covered patients with inherited bone marrow failure syndromes (Fanconi anemia, SBDS), pregnant or lactating females, secondary AML, acute promyelocytic leukemia, Philadelphia chromosome positive AML, mixed-phenotype or bilineal leukemia, JMML, and t-AML. Diagnostic RNA-sequencing data was available for a total of 1,410 individuals (dbGaP Study Accession: phs002187.v1.p1). Consent was obtained from all study participants in accordance with the Declaration of Helsinki. The Fred Hutchinson Cancer Center IRB and the COG Myeloid Biology Committee approved and oversaw the use of patient data.

### Non-negative Matrix Factorization and Gene Set Enrichment Analysis

Non-negative Matrix Factorization^56,57^ (NMF) was performed on the log-normalized RNA sequencing results using the Rcpp Machine Learning library (github.com/zdebruine/RcppML), with Monte Carlo cross-validation on 3 random replicates using Wald’s method to identify an optimal rank near k = 100. The sample loadings in the “H” matrix were used for UMAP projection and for enrichment analysis of factors within *FUS::ERG* positive samples. Gene set enrichment analysis was performed using the R package msigdbr (Dogalev, 2022) on the gene weights within specific factors from NMF of TARGET AML patient RNAseq data by way of RcppML^58,59^. NMF factors of interest were selected based on enrichment or depletion in *FUS::ERG* positive cases, and positively enriched gene sets were ranked by FDR adjusted p-value. The top 20 MSigDB GO:BP gene sets were visualized using the R-package ggplot2^60^.

### H3K27ac chromatin immunoprecipitation–sequencing processing

Data provided from pAML patients at Texas Children’s Hospital. Methods are as previously published.^26^

### Cell culture

TSU-1621MT cells (RRID:CVCL_A455) were generously provided by Dr. Olaf Heidenreich of the Princess Máxima Center, Utrecht, The Netherlands. YNH-1 (ACC 692) and OCI-AML3 (ACC 582) cells were purchased from the Leibniz Institute DSMZ-German Collection of Microorganisms and Cell Cultures. Cells were cultured at 37ᵒC in 5% CO2 in RPMI Medium 1640 (Life Technologies, Grand Island, NY) supplemented with 15% heat-inactivated FBS (Life Technologies), 56 U/ml/56 μg/ml penicillin/streptomycin (ThermoFisher Scientific, Waltham, MA) and 10 ng/ml human recombinant G-CSF (Stemcell Technologies, Vancouver, Canada).

### Reagents

Tazemetostat (EPZ-6438) was purchased from Selleck Chemicals (Houston, TX). Recombinant human IFN-γ was purchased from R&D Systems (Minneapolis, MN). H3K27me3 (C36B11) Rabbit mAb #9733 was purchased from Cell Signaling Technology (Danvers, MA, USA). MS177 PROTAC was purchased from MedChemExpress (Monmouth Junction, NJ, USA). Leukocyte reduction cones were purchased from Versiti (Milwaukee, WI, USA). Human IL-2 recombinant protein was purchased from Thermo Fisher. RBC Lysis Buffer (10X) was purchased from Biolegend (San Diego, CA, USA).

### Flow cytometry

For MHC I and II analysis, cells were incubated with anti-human HLA-A, B, C antibody conjugated to Pacific Blue^TM^ (Biolegend) and anti-human HLA-DR, DP, DQ antibody conjugated to Alexa Fluor 647 (Biolegend). Dead cells were labeled with Propidium Iodide Ready Flow^TM^ Reagent (ThermoFisher Scientific). For immune coculture experiments, TSU-1621MT cells were labeled with CellTrace™ Violet Cell Proliferation Kit (ThermoFisher Scientific) and quantified using CountBright™ Plus Absolute Counting Beads (ThermoFisher Scientific). Samples were processed in the VAI Flow Cytometry Core (RRID:SCR_022685) on a Beckman Coulter CytoFLEX S cytometer and analyzed using FlowJo (RRID:SCR_008520) software (BD Life Sciences).

### Statistical analysis

For all analysis, a p value of ≤ 0.05 was considered statistically significant. Mantel-Cox Log-rank tests were used to analyze survival curves in Figure 1 and 7. An unpaired t test was performed for analyses between two groups; for analyses involving multiple groups, data was analyzed by one-way ANOVA followed by Šídák’s multiple comparisons tests. A two-proportion z-test was used to analyze Figure 1D and 7C. Pearson correlation coefficients were calculated for scatterplots in Figures 3 and 5. A one-sample t test was used to analyze normalized data in Figure 3D.

## Supplemental Material

**Supplemental Table 1.** Gene weights for NMF factors. (See supplemental CSV file.)

**Supplemental Figure 1.**
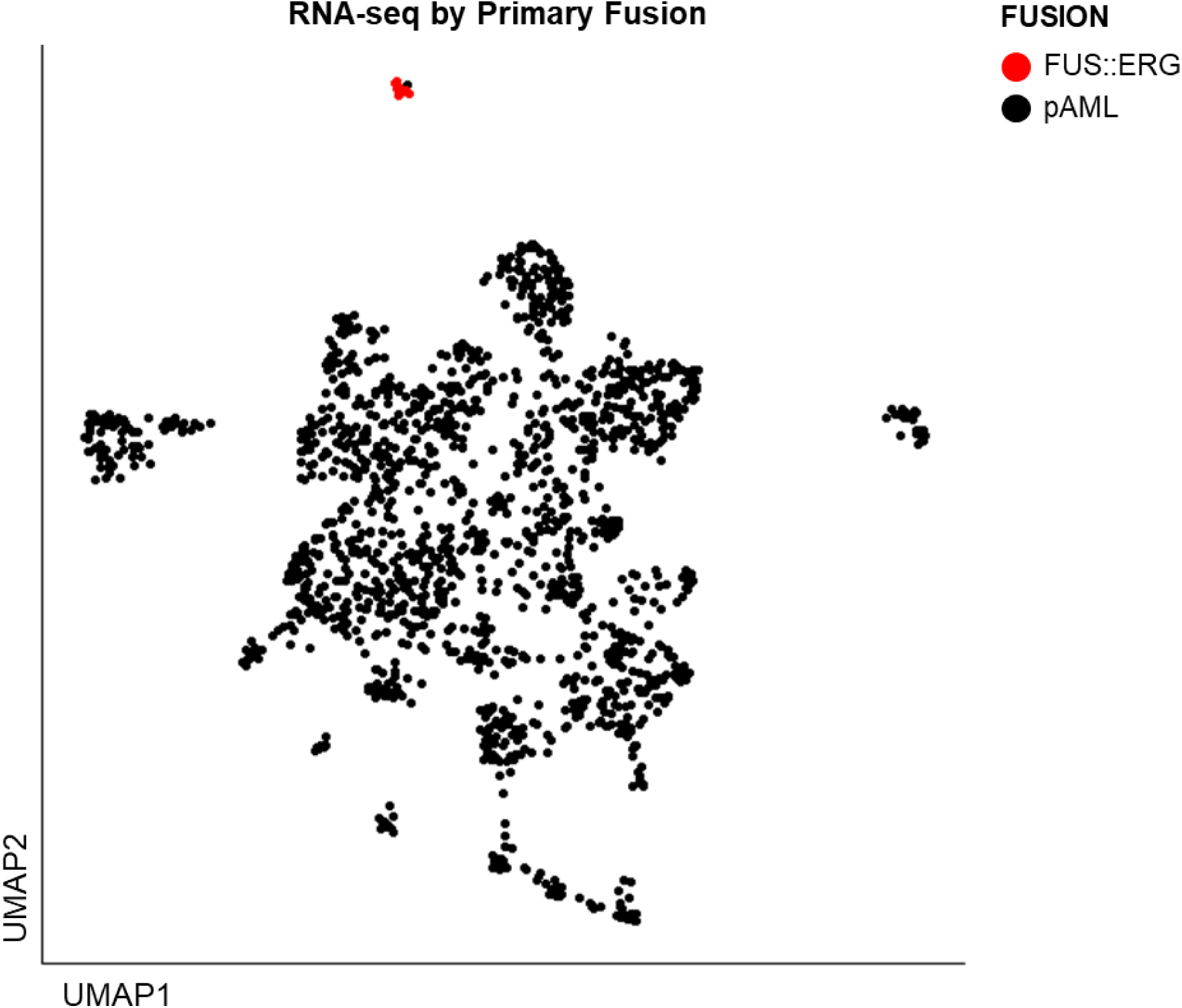
UMAP projection of gene expression from 1410 AAML0531/1031 patients who had RNA-sequencing at diagnosis. *FUS::ERG* patients highlighted in red.

**Supplemental Figure 2.**
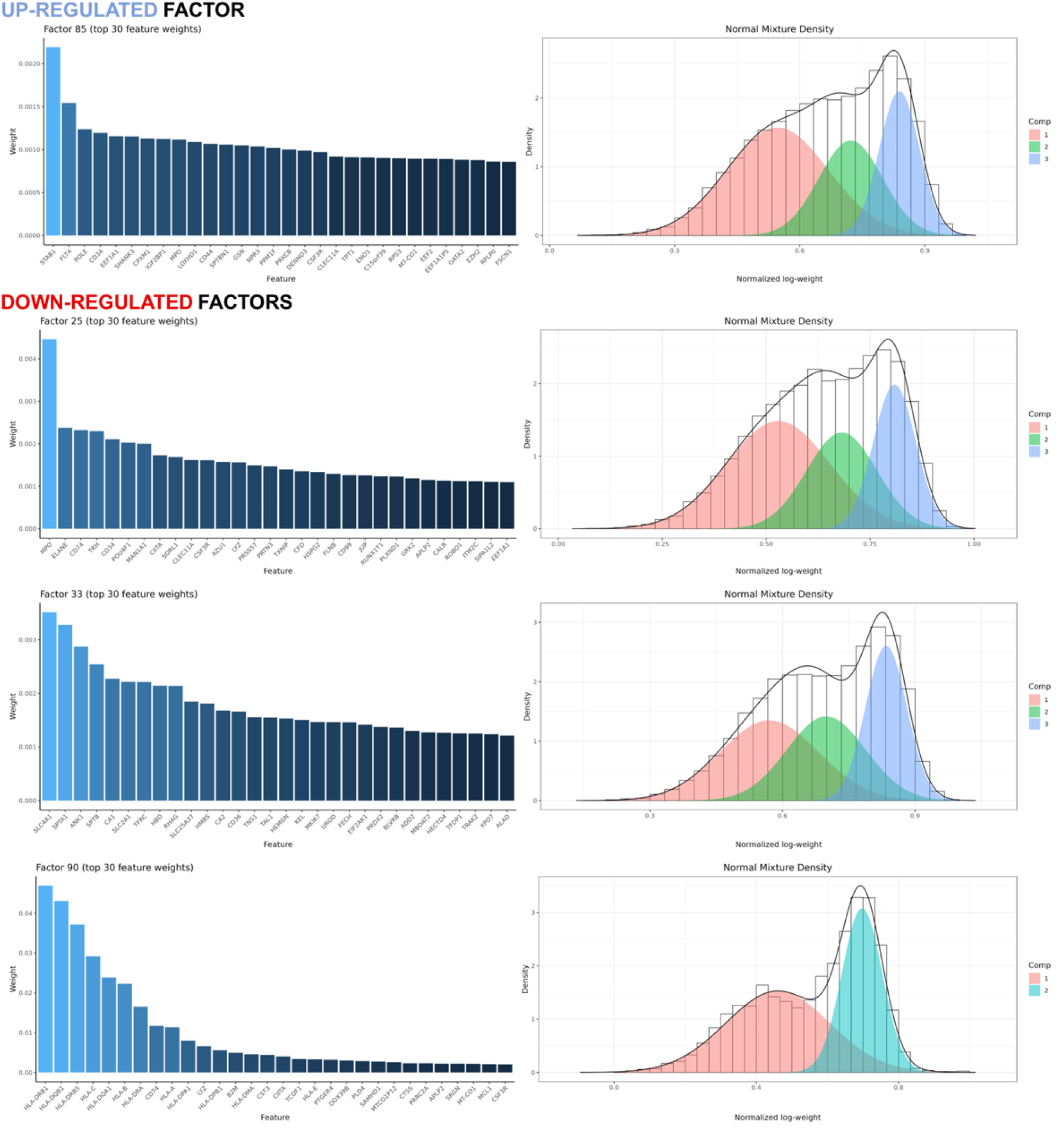
Top 30 feature weights for NMF factors 25, 33, 85, and 90.

**Supplemental Figure 3.**
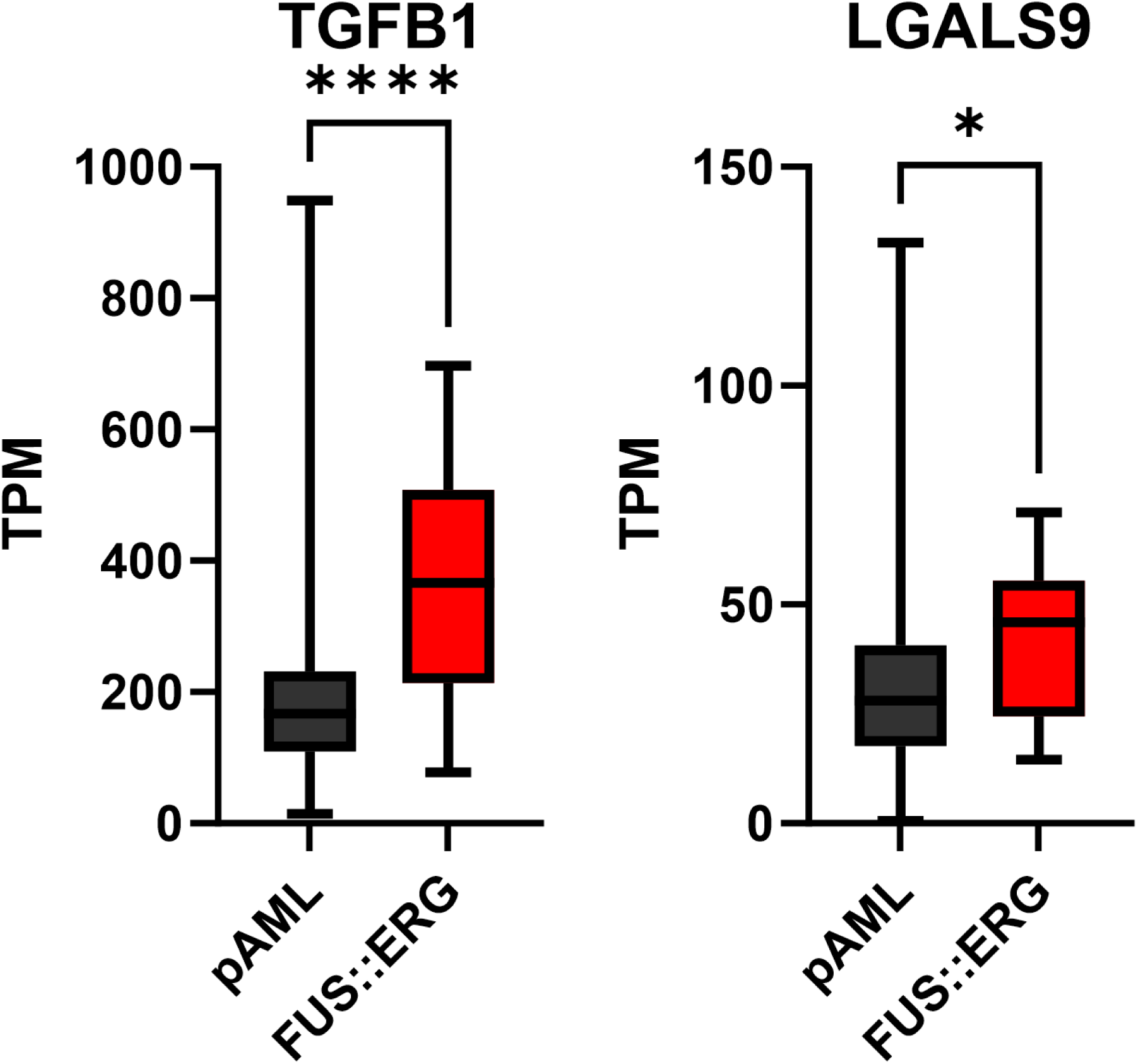
Expression of pro-Treg stimulants *TGFB1* and *LGALS9* in pAML versus *FUS::ERG* (n = 1410). ****p-val ≤0.0001; *p-val ≤0.05.

**Supplemental Figure 4.**
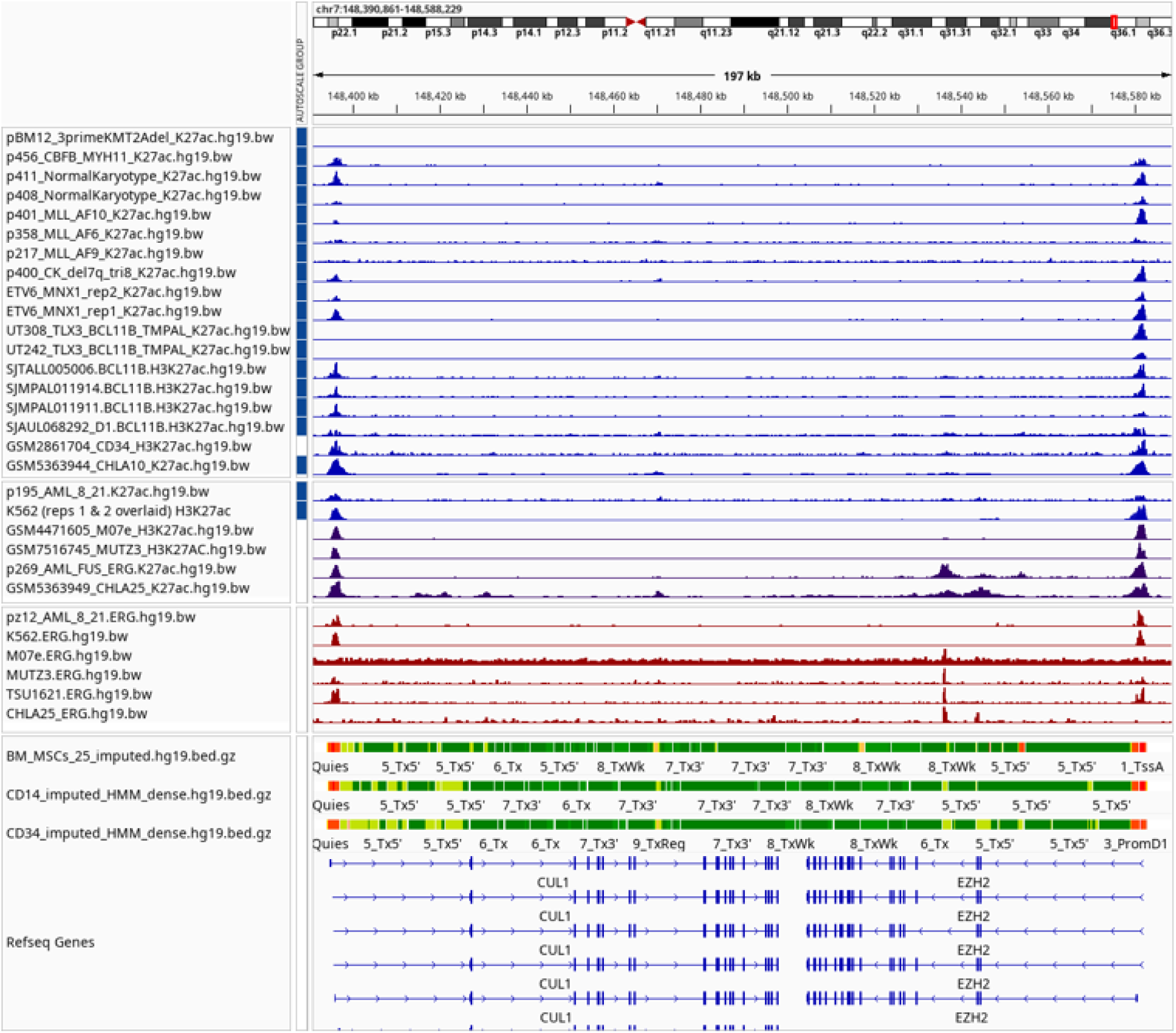
H3K27ac and ERG ChIP-seq^25,61,65–70^.

**Supplemental Figure 5.**
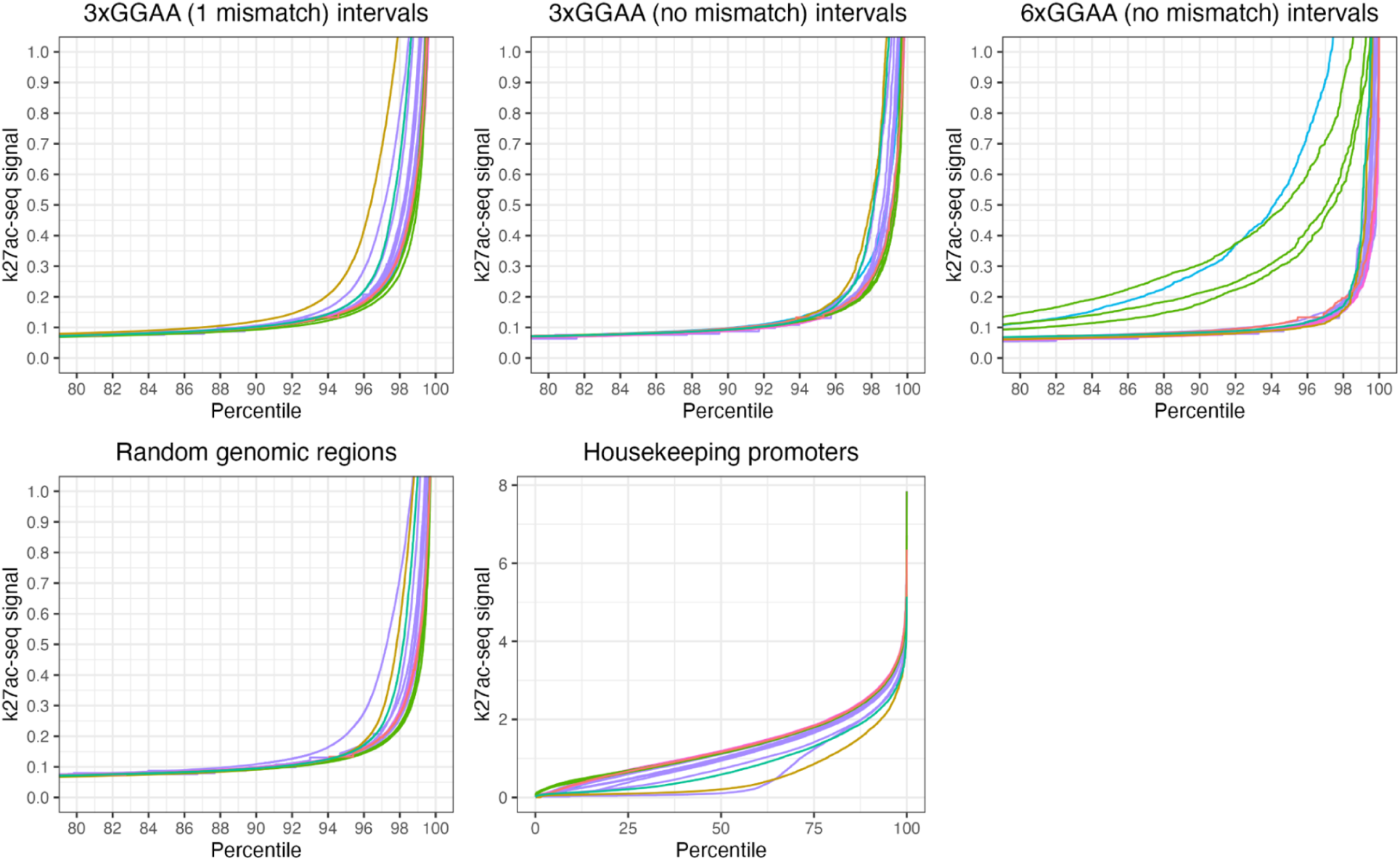
H3K27Ac analysis at 6x GGAA motifs conducted on primary pAML^26^, T-lymphoid/myeloid mixed-phenotype acute leukemia (TMPAL)^61^, Desmoplastic small round cell tumors (DSRCT)^62^, and Ewing’s Sarcoma^63^ patients with CD34 hematopoietic progenitor cells (HPC) and mesenchymal stem cell (MSC) controls^64^; pAML *FUS::ERG*^26^ cases highlighted in blue. Analysis was conducted on 3x repeats with one mismatch (A), 3x repeats with no mismatch (B), and 6x repeats with no mismatch (C). Random genomic regions and housekeeping promoters shown in (D) and (E) as controls.

